# Polygenic prediction of the phenome, across ancestry, in emerging adulthood

**DOI:** 10.1101/124651

**Authors:** Anna R. Docherty, Arden Moscati, Danielle Dick, Jeanne E. Savage, Jessica E. Salvatore, Megan Cooke, Fazil Aliev, Ashlee A. Moore, Alexis C. Edwards, Brien P. Riley, Daniel E. Adkins, Roseann Peterson, Bradley T. Webb, Silviu A. Bacanu, Kenneth S. Kendler

**Author notes:** Both authors contributed equally to the manuscript (Anna R. Docherty, Arden Moscati). Corresponding author: Departments of Psychiatry and Human Genetics, University of Utah School of Medicine, 501 Chipeta Way, Salt Lake City, Utah, USA.

## Abstract

**Background:** Identifying genetic relationships between complex traits in emerging adulthood can provide useful etiological insights into risk for psychopathology. College-age individuals are under-represented in genomic analyses thus far, and the majority of work has focused on clinical disorder or cognitive abilities rather than normal-range behavioral outcomes.

**Methods:** This study examined a sample of emerging adults 18-22 years of age (N = 5,947) to construct an atlas of polygenic risk for 33 traits predicting relevant phenotypic outcomes. Twenty-eight hypotheses were tested based on the previous literature on samples of European ancestry, and the availability of rich assessment data allowed for polygenic predictions across 55 psychological and medical phenotypes.

**Results:** Polygenic risk for schizophrenia in emerging adults predicted anxiety, depression, nicotine use, trauma, and family history of psychological disorders. Polygenic risk for neuroticism predicted anxiety, depression, phobia, panic, neuroticism, and was correlated with polygenic risk for cardiovascular disease.

**Conclusions:** These results demonstrate the extensive impact of genetic risk for schizophrenia, neuroticism, and major depression on a range of health outcomes in early adulthood. Minimal cross-ancestry replication of these phenomic patterns of polygenic influence underscores the need for more genome-wide association studies of non-European populations.

## Introduction

Broad phenotyping can greatly enhance our understanding of the underlying structure of genetic vulnerability to psychiatric disorders. Thus, genome-wide polygenic risk research is increasingly considering batteries of clinical phenotypes in “phenome-wide” studies (Bulik-Sullivan *et al.* 2015; Krapohl *et al.* 2015; Hagenaars *et al.* 2016). One promising approach emerging from the phenome-wide genomic literature uses published summary statistics from large genome-wide association studies to calculate genome-wide polygenic scores (GPS) for an array of major disorders and clinically relevant traits. These scores are then used to predict a number of potentially informative psychiatric, psychological and physical health phenotypes. Along with cross disorder genomic research examining the co-heritability of major psychiatric disorders (e.g., LD regression) (Cross Disorder Group of the Psychiatric Genomics Consortium, 2013), polygenic phenomic approaches are among the most promising methods for elucidating the complex overlapping genetic architecture of psychiatric disorders and discovering unexpected genotype-phenotype associations (Docherty et al. 2016). However, previous research has suffered from a restricted range of phenotypes, and has not included, for instance, GPS of anxiety, eating, and inflammatory disorders, personality, lipid levels and puberty traits in the prediction of outcome phenotypes (which have generally focused on cognitive abilities) and has not examined samples between the ages of 17 and 35, the developmentally critical stage encompassing emerging and young adulthood.

This study applied such an approach to GPS (33 total) in a phenotypically extensive genetic study of emerging adulthood outcomes. Emerging adulthood, a period starting at the age of 18 when adolescents begin to develop the roles and independence of adulthood, reflects a high-risk age range for the onset of many psychiatric and substance use disorders, including schizophrenia, affective disorders, anxiety disorders and alcohol and drug use disorders. Data from the National Comorbidity Survey Replication sample indicate that three quarters of all lifetime cases of DSM-IV diagnoses start by age 24 (Kessler *et al.* 2005), and WHO’s World Mental Health data indicates that approximately three quarters of lifetime psychiatric disorders begin by the mid-20′s (Kessler *et al.* 2007).

The University Student Survey (called “Spit for Science”, or S4S) was developed to identify risk factors for onset of mental health disorders with large-scale assessment of genetic, environmental, and developmental influences. Discovery summary statistics from 33 genome-wide association studies (GWAS) were used to derive GPSs in this large sample of young adults (N = 5,947) across a range of psychiatric, psychological, and physical health traits (Table S1 in online supplementary materials). Expanding on previous research, twenty-eight hypotheses of genetic prediction were tested based on selected studies in past literature. Further, the availability of rich clinical assessment data allowed for the calculation of polygenic predictions across a greater number of outcomes than has ever been studied previously, many of which were completely novel in phenomic studies. These included 55 psychiatric, psychological and medical phenotypes (listed in Table S2 in the online supplementary material).

Moreover, the GPS metrics were powerful enough to examine relationships across subsamples of different ancestries. While GWAS approaches require tens of thousands of individuals to locate “hits,” continuous polygenic scores require far smaller samples for adequate power. This sample was suitably diverse in ancestry to map the GPS-phenotype in young adults of European ancestry (EUR, N=3,016) and then to replicate these findings across non-European ancestry groups including of African origin (AFR, N=1,339), native American origin (AMR, N=581), and East Asian (EAS, N=557), and South Asian origin (SAS, N=454). Separate association matrices were created for the empirically categorized AFR, AMR, EAS, and SAS samples and are provided here and in the supplemental figures available online.

We can learn much from the study of emerging adults over and above adolescent samples, as early behavioral patterns that may precede adult psychopathology can be studied, and new hypotheses about critical exposures and environmental risk factors can emerge. The results presented here reflect a polygenic modeling framework in a large young adult sample, and provides evidence that the integration of phenotypic and genotypic data will be useful in the prediction of negative health outcomes in emerging adults.

## Methods

### Sample Ascertainment and Phenotyping

Phenome-wide behavioral data (N=7,592) were drawn from young adults from the first three cohorts in S4S, samples drawn from a large urban university in the Mid-Atlantic United States, which included 5,947 unrelated individuals with genome-wide genotypes (Dick *et al.* 2014). The S4S sample does not overlap with any of the discovery GWAS samples used in these analyses. Details of participant ascertainment have been published elsewhere (Dick *et al.* 2014) but briefly, emerging adults ages 18-22 were recruited across multiple cohorts, for a campus-wide study of genetic and environmental factors contributing to alcohol and substance use. The protocol was approved by the university Institutional Review Board, and carried out in accordance with the provisions of the World Medical Association Declaration of Helsinki. Participants were 61.1% female with a mean age of 18.59 at first assessment. Representativeness of this sample is strong and has been reported elsewhere (Dick *et al.* 2014). Assignment to ancestry group was empirically based on greatest similarity to 1000 Genomes Phase 3 superpopulations. The present analyses included 55 traits from the domains of psychopathology, personality, health factors, and educational achievement (Table S2 in the online supplementary materials). All analyses included age, sex, and 10 ancestry principal components as covariates. Variables assessed at multiple occasions or in multiple cohorts were adjusted for number of assessments and cohort group. Sample sizes for each of the phenotypic measures are also provided in Table S2.

### Genetic Risk Scoring

DNA collection, calling, and imputation is detailed elsewhere (Dick *et al.* 2014). We processed genotypes using standard quality control procedures followed by imputation of SNPs using the 1000 Genomes Project reference panel. After imputation and quality control, we included approximately 2.3 million variants into the polygenic scoring analyses. A GPS for each discovery phenotype was calculated using the summary statistics we obtained from 33 GWAS (Table S1 in the online supplementary materials). Python-based LDpred (Vilhjálmsson *et al.* 2015) was used for these analyses because of its ability to account for linkage disequilibrium (LD) structure (Krapohl *et al.* 2015) using our own large EUR test sample, and its use of all genetic variants (instead of specified p-value threshold for inclusion of the genetic variants in the GPS). LDpred allows for the modeling of LD based on LD in the discovery sample to weight the relative contributions of syntenic variants to the outcome phenotype. LDpred uses postulated proportions of causal variants in the genome as Bayesian prior probabilities for GPS calculations, and we tested a range of different priors (proportions of 0.3, 0.1, 0.03, 0.01, 0.003, and 0.001), as well as the model of infinite variants of infinitesimally small effect (Fisher, 1919) to construct scores.

### Phenotype Prediction

A flowchart depicting the GPS-phenome cross-ancestry prediction and GPS-GPS correlation procedure is presented in Figure 1. Regressions were run using R to compare full (GPS, ten ancestry principal components, age, sex, cohort, and number of measurements when applicable) and restricted models where GPS was removed. Prior to the global analyses, a set of *a priori* hypotheses, gathered from previous research, were tested (Table 1). We elected to generate several hypotheses prior to analysis, because some literature was available to support previous evidence of relationships between GPS and outcome. We elected to forgo experimental binning (into quantiles, for example) in order to minimize the number of exploratory analyses beyond regressions of GPS on the phenotypes. Multiple testing was corrected for using a False Discovery Rate (FDR) of 5% (Benjamini & Hochberg, 1995) within each ancestry group using the p.adjust function in R; the FDR is appropriate for an analysis designed to evaluate the pattern of relationships between many constructs because it treats each combination of discovery phenotype, outcome, and LDpred prior level as an independent test, despite the presence of positive dependency between many of these tests. It should be noted that this multiple testing correction did not account for previously established associations or for the correlations observed in our sample, between multiple prior levels tested in the same discovery phenotypes. This was an added attempt to filter out potentially spurious results.

### Cross-Disorder GPS Partial Correlations and GPS-GPS Replication Hypotheses Across Ancestry

In addition to testing the GPSs prediction of the phenotypes, GPSs were also examined for correlations with each other in all ethnicities. These provide different results than genetic correlation estimates, but are intended to demonstrate that GPS scores are not independent, and that variance attributable to a particular discovery phenotype may be partially shared with another. This sharing may be due to common genetic factors between phenotypes, possible sample overlap, and error variance. GPS correlations have been previously reported in EUR, but this analysis added phenotypes such as cardiovascular and triglyceride factors. Correlation coefficients, *p*-values, and *q*-values (after correcting the p-values for the FDR of 5%) were derived for GPS partial correlations using R and adjusting for the ancestry principal components. We chose to use partial correlations in order to standardize the weights across phenotype and provide more direct comparisons of statistics for plotting purposes. Based on the cross-disorder psychiatric genomics findings to date (Bulik-Sullivan *et al.* 2015), we expected significant GPS associations between schizophrenia (SZ) and bipolar disorder (BP), SZ and autism (AUT), SZ and major depressive disorder (MDD), BP and MDD, and AUT and attention deficit hyperactivity disorder (ADHD) across each of the ancestry groups (see Table 2).

## Results

### Genetic Profile Score-Phenotype Prediction

#### *A Priori* Replication Analyses

We evaluated previous cross-phenotype predictions based on recent work—for example, that age at menarche had an inverse association with obesity/body mass index (Bulik-Sullivan *et al.* 2015). We tested several hypotheses in the European group, in order to maximize sample size without introducing potential population stratification. The multiple testing correction procedure we chose (FDR) was suitable for these analyses, given the positive dependency between many of the tests, allowing us to correct uniformly for the total number of tests while still keeping type I error rate relatively controlled. Of the 28 predictions tested, 22 showed effects in the expected direction (p=0.002, one-tailed sign test), and 7 were significant after stringent multiple-testing correction. Two previous notable null associations, MDD GPS predicting Grade Point Average (GPA), and Type 2 Diabetes GPS predicting GPA, were also null in our sample. Full results are presented in Table 1, including additional associations with the listed GPS phenotypes.

#### Phenome-Wide Prediction

We also performed hypothesis-free analyses across all 33 GPS and 55 S4S phenotypes to explore potentially novel associations. Multiple prior proportions of causal variants in the genome were tested, as detailed in Methods. Figure 2 presents notable results for GPS prediction of phenotypes in the European group for the prior proportion of 0.3 (that is, an initial assumption that 30% of the genome is associated with the GPS phenotype). The 0.3 prior level showed stronger prediction in past work (Krapohl *et al.* 2015), and corresponds to a plausible assumption about the genetic architecture of many complex traits, due to instances of increasing sample size of GWAS proportionally increasing numbers of associated loci. In this group and prior proportion level, out of 1,815 associations 35 were between *q* < 0.16 and *q* > 0.05 (0.16 being the P-value threshold corresponding to Akaike Information Criterion (Akaike, 1974)), 11 between *q* < 0.05 and *q* > 0.01, and 26 *q* < 0.01. An additional 53 associations showed at least suggestive significance at other prior levels. A heatmap of analyses at can be found in Figure 2 (EUR; and for replications in all ancestries, Figures S1-S4 available in the online supplementary materials). Each plot presents significant associations as well as the direction of effect. Because of the uniform correction for multiple testing, we included of interest q<0.16 associations, which would be significant with more traditional correction methods accounting for previously established associations.

Notable results included SZ GPS significantly predicting nicotine use, depression and anxiety symptoms, and family history of depression, anxiety, alcohol use disorder, and drug use. In addition, GPS for neuroticism (N) predicted a number of relevant psychiatric phenotypes, including neuroticism, depression and anxiety symptoms.

### Genetic Profile Score Prediction of the Phenome Across Non-European Ancestries

As noted earlier, most discovery GWAS have used European samples, and while there is good evidence for cross-ancestral replication for some traits, the generalizability of many of these relationships across ancestry is not known. The diverse ancestry groups within S4S allowed cross-ancestral replication, and the use of continuous GPS metrics made the sample sizes available powerful enough to examine these hypotheses. A small proportion of the strongest predictors observed in the EUR were replicated across the other ancestries, with a broadly similar pattern of results across all ancestry groups only for “basic” traits such as height and BMI. While some outcome phenotypes were strongly predicted by GPS, a few outcome phenotypes, including physical activity, lifetime history of panic attack, age at first sexual intercourse, and bulimia nervosa were not predicted by any GPS in any ancestry group. In addition, some expected associations (e.g., PRS for nicotine use predicting smoking behaviors) while in the expected direction, were not significant in this sample.

Most associations of SZ GPS with outcome traits observed in EUR did not reach significance in other ancestry samples. In addition, some novel associations were observed in other ancestries. For example, lifetime smoking GPS was positively associated with number of alcoholic drinks per day in AMR. Neuroticism GPS was positively associated with stressful life events, trauma (interpersonal and general), and PTSD in SAS. These patterns of effects are based on EUR GWAS summary statistics, and must be replicated using AMR and SAS GWAS summary statistics in the future. However, they suggest potential pleiotropic effects relevant to outcomes in these populations.

### GPS-GPS Correlations

#### *A Priori* Hypothesis Testing and Global Cross-Disorder Genetic Profile Analyses

*A priori* hypotheses (described in the Methods and listed in Table 2) of relationships between GPS scores were tested at a prior proportion level of 0.3. Figure 3 presents the results for GPS-GPS partial correlations at a GPS *p* = 0.3, and these results are presented because some phenotypes studied here (e.g., Neuroticism) were not included in previous analyses. Notable unexpected correlations were observed, including significant positive correlations of neuroticism GPS with GPSs for triglycerides and coronary artery disease. These associations also serve as evidence of non-independence across traits in this sample. Finally, Figures S5-S8 (available in the online supplementary materials) present these same correlations across the four non-EUR ancestry groups. There is some overlap between the discovery samples for neuroticism and triglycerides, but no overlapping studies were included in the neuroticism and the coronary artery disease discovery samples. Therefore, the correlation of neuroticism and coronary artery disease is especially likely to reflect underlying genetic correlation between neuroticism and artery disease. Despite overlap in the discovery samples for the neuroticism and triglycerides polygenic scores, validation using LD score regression supported the existence of a genetic relationship between them (*r*_*G*_ = 0.53; *SE* = 0.04; *p* = 1.5 × 10^−36^).

## Discussion

The findings here present a wide-ranging and nuanced picture of major dimensions of vulnerability to psychopathology at a genetic level. This study includes substantial sample sizes of emerging adults, uses outcome measures (with novel phenotypes in phenomic analyses; see Table S2 in the online supplement for details of assessment scales), includes a wide range of discovery GWAS, and is powerful enough to draw preliminary conclusions about several ancestries. Because this study does not look for “hits” in the traditional GWAS sense and instead uses continuous GPS metrics, sample sizes provide adequate power across all separate ancestries in this study.

Importantly, results reflect EUR relationships between anxiety, depressive, and schizophrenia-spectrum disorders that are largely consistent with current conceptualizations of diagnostic classification, and confirm the important involvement of a network of medical and risk phenotypes in genetic predisposition to these disorders. Informative genetic associations between medical and clinical phenotypes exist despite the relative dearth of individual loci of genome-wide significance.

We can learn a lot from the study of emerging adults relative to younger, adolescent samples, as more targeted theories about critical exposures and environmental risk factors can emerge. For example, GPS for SZ predicted anxiety, depression, nicotine use, experiences of interpersonal trauma, and family history of mental health problems. Importantly, these results expand on recent evidence that genetic risk for SZ can successfully predict diverse risk phenotypes such as anxiety and negative symptoms (Kendler *et al.* 1996; Fanous *et al.* 2001; Docherty & Sponheim, 2008; Docherty & Sponheim, 2014; Docherty *et al.* 2015; Jones *et al.* 2016; Kendler, 2016), and demonstrate important links between SZ genetic risk and health factors in early adulthood. Significant association of GPS with easily measured, specific risk factors (e.g., nicotine use, family history, trauma) indicates that GPS could be useful in predicting psychopathology, particularly in conjunction with environmental moderators.

The incorporation of personality traits such as neuroticism was also quite informative. For example, neuroticism GPS significantly predicted a broad network of general anxiety, phobia, panic, neuroticism, and depression phenotypes in EUR, as well as multiple health-related GPSs. This is consistent with previous biometrical and genomic research reporting significant relationships of neuroticism with MDD (Kendler & Myers, 2010; Genetics of Personality Consortium *et al.* 2015; Docherty *et al.* 2016), and preliminary findings from the UKBiobank suggesting a genetic overlap of neuroticism with cardiovascular health (Gale *et al.* 2016). Conversely, GPS for extraversion predicted fewer depressive symptoms, fewer anxiety symptoms, and less family history of mental health problems, though these associations did not remain significant after multiple testing correction. Associations pertaining to GPS for well-being in this sample are forthcoming from our research group.

Notable unexpected GPS-GPS results included positive correlations of neuroticism GPS with GPSs for coronary artery disease, which is likely to reflect underlying genetic correlation, as well as with triglycerides. This is the first study we know of to document significant positive genetic associations between neuroticism and cardiac health, despite the high public health cost of neuroticism being well-documented (Cuijpers *et al.* 2010; cardiovascular risk and association with psychiatric phenotypes like neuroticism may be of special interest to public health efforts). Most of the GPS-GPS *a priori* relationships chosen for replication testing were represented in the same direction across all ancestry groups, corroborating previous efforts to map relationships between genetic risk profiles.

The abundance of significant relationships between intuitive combinations of GPSs and related outcomes is reassuring considering the many factors that could attenuate the statistical link between them. Association between a GPS and an outcome not only reflects correlation between the phenotype in the original (‘discovery’) GWAS that produced the statistics used to compute the GPS and the outcome phenotype, but is also related to a number of other factors. The link is limited by how accurately the GWAS measured the initial phenotype, how similar the discovery and test samples are (in age, ancestry composition, proportions of each sex, etc.), how well the test phenotype is measured by the data collection instrument, and how well it can incorporate indirect pathways from the genetic architectures to either phenotype.

For example, physical activity increases HDL levels (Kokkinos & Fernhall, 1999), so those who had higher HDL levels in the discovery GWAS (Teslovich *et al.* 2010) were likely a mix of those with innately high levels, those who engaged in higher levels of physical activity, and those with both traits. Therefore, HDL GPS perhaps indexes some propensity to engage in HDL-promoting behaviors, in addition to HDL metabolic variation such as a slower rate of HDL catabolism, which is thought to be the most common genetically determined mechanism of increased HDL levels in humans (Rader, 2006). The portion of the HDL GPS due to fitness behaviors may explain some of the polygenic association with the test phenotypes of BMI and weight. Of note, while the HDL GPS did not significantly predict the physical activity phenotype in S4S, the direction of effect was positive, and that particular phenotype had one of the smallest sample sizes, at 43 3 EUR individuals, and therefore lower power than others.

Using EUR GWAS summary statistics produced differential relationships of GPS with outcomes across ancestry, with few effects replicating across ancestry groups. This could be due to decreased power to detect effects given the smaller sample sizes of the other ancestry groups. However, within each ancestry group, new significant effects across GPS and outcome measures were observed as well. These effects suggest ancestry-specific summary statistics, or those with larger samples, may reveal differences in the pattern of relationships between phenotypes in different groups. Results underscore a need for more GWAS of non-EUR ancestry samples.

There are a number of limitations to be aware of when interpreting these results. The synthesis of information from so many sources compounds any methodological and psychometric issues present in the original studies, so there is probable bias in multiple levels of the analysis that is difficult to measure. It is unclear how generalizable our results are to the general population. However, we might assume that bias in the college sample would cause less robust associations with psychopathology, and that effects might be more pronounced in the general population. In order to maintain proximity with real outcomes, we did not transform our phenotypic variables to increase normality, but standardized the continuous variables computed from the participant responses to maintain comparable ranges of measurement. However, none of the phenotypes in which we found significant results evidenced high skew or kurtosis, so it is unlikely that significant effects were due to non-normal phenotype distributions. While LDpred performs adequately across ancestry groups, the accuracy in non-European ancestry groups is attenuated to the degree that multiple causal variants fall in regions where LD patterns differ across ancestry. In addition, a recent pre-print (Martin *et al.* 2016) shows biased predictions in several different populations using GPS for phenotypes that are also used in this paper (for example, Type 2 diabetes and SZ). Finally, we observed some differential effects across priors in our analyses. Until these effects are replicated at a given prior, or there is justified precedent in the literature, we are unable to choose one ideal prior.

Overall, this broader picture of genetic vulnerability has important implications for how we study risk and resilience in emerging adulthood. While the variance explained by any of these GPSs is small, (for instance, the largest R^2^ for predicting the height phenotype was 0.055, from the height GPS at a prior level of 1) they provide easily accessible information to guide future prediction, prevention, and intervention efforts to improve health and quality of life outcomes. Future longitudinal and intervention research could elaborate on this atlas to examine the predictive validity and prevention utility of many of the phenotypes here, such as neuroticism, family history, trauma, and nicotine use. This research also suggests that future polygenic work would benefit from GPSs based on non-European ancestry groups when such summary statistics are available. Phenome-wide research utilizing deeper phenotyping methods will likely further enhance results, and thus future prediction of positive and negative health outcomes.

Finally, the relationships outlined here provide implicit suggestions for studies of the causal structure of the GPS phenotypes themselves. The genetic architecture of most of the traits and disorders in the atlas display substantial overlap; a significant portion of genetic variation involved in the etiology of these constructs does not selectively contribute to risk for one phenotype as we know it, but rather has effects that act on some axis of liability that increases the likelihood of many phenotypes. Analyzing multiple related phenotypes in a holistic fashion allows elucidation of the individual patterns of genetic and environmental factors that may explain causal mechanisms— which risk factors they share, and which are unique to one phenotype, thus serving to refine our nosological theories. Any epidemiological analysis is limited if the construct under study is not a uniform disease entity, but as characterization of constructs improves, the power to find their correlates does as well. The better we ask the questions, the more useful the answers become, for both clinical and scientific purposes.

## Author Contributions

R. Docherty, A. Moscati, and K. S. Kendler developed the study concept. A. Moscati, J. E. Savage, J. E. Salvatore, and M. Cooke contributed to psychometric analyses and data collection. D. Dick and K. S. Kendler oversaw data collection. A. Moscati and A. R. Docherty performed the data analysis and interpretation under the supervision of B. T. Webb, S. A. Bacanu, D. E. Adkins, F. Aliev, A. C. Edwards, and B. P. Riley. A. R. Docherty and A. Moscati drafted the manuscript, and K. Kendler, A. C. Edwards, J. E. Savage, J. E., Salvatore, M. Cooke, B. T. Webb, S. A. Bacanu, D. E. Adkins, A. Moore, R. Peterson, and D. Dick provided edits. All authors approved the final version of the manuscript for submission.

## Acknowledgements

We would like to thank the participants and the many VCU faculty, students, and staff who contributed to the design and implementation of this project.

## Financial Disclosures

All authors declare no conflict of interest with respect to the authorship or publication of this article. Data collection for the study was funded by R37AA011408, P20AA107828, K02AA018755, and P50AA022537 from the National Institute on Alcohol Abuse and Alcoholism, by Virginia Commonwealth University, and by UL1RR031990 from the National Center for Research Resources and National Institutes of Health Roadmap for Medical Research. A. Docherty was funded by K01MH109765 from the National Institute of Mental Health, and by a Brain & Behavior Research Foundation (formerly NARSAD) Young Investigator Award. A. Moscati, A. Moore were supported by institutional training grant T32MH20030. J. Salvatore was supported by F32AA22269 and K01AA024152. M. Cooke received support from UL1TR000058 from the National Institutes of Health National Center for Advancing Translational Science. A. Edwards was supported by K01AA021399.

## Declaration of Interest

Authors report no conflicts of interest.

